# Non-Redundant Roles of Topoisomerase 2α and 2β in the Cytosolic Replication of Vaccinia Virus

**DOI:** 10.1101/2025.02.05.636656

**Authors:** Ilaria Dalla Rosa, Lois Kent, Michael Way

**Affiliations:** Cellular Signalling and Cytoskeletal Function Laboratory, The Francis Crick Institute, 1 Midland Road, London, NW1 1AT, UK; Proteomics Science Technology Platform, The Francis Crick Institute, 1 Midland Road, London, NW1 1AT, UK; Department of Infectious Disease, Imperial College, London W2 1PG, UK

## Abstract

Vaccinia virus is a large enveloped DNA virus, which, like all poxviruses, replicates in the cytoplasm of infected cells. Vaccinia was historically thought to encode all the proteins required for its replication. However, more recent findings have shown that nuclear host proteins are redirected to the cytoplasm to facilitate viral replication. Among these, topoisomerase 2α (TOP2A) and 2β (TOP2B), which mediate nuclear transcription, DNA replication, and chromosome segregation are the most abundant host proteins associated with nascent viral genomes. Here, we investigate the mechanisms driving TOP2A and TOP2B cytoplasmic translocation and their role in viral replication. We found that early viral protein synthesis induces the cytosolic relocalization of both isoforms, which are subsequently recruited to viral factories by an interaction of their C-terminal domains with the viral ligase, A50. TOP2A promotes replication by interacting with the vaccinia DNA replication machinery. In contrast, TOP2B suppresses replication by enhancing the formation of double-stranded RNA and antiviral granules, containing components of the tRNA splicing ligase complex. Our analysis provides new insights into host-pathogen interactions during poxvirus infection and the role of topoisomerase 2 outside of the nucleus.

## Introduction

Poxviruses are a family of large enveloped DNA viruses characterized by a unique replication cycle that occurs entirely in the cytoplasm of infected cells (1,2). Notable members of the family include the causative agents of smallpox and mpox (formerly known as monkeypox) (3). Vaccinia virus, the most studied family member is regarded as the prototypical poxvirus, which was used as the vaccine in the global eradication of smallpox (4). The large, double-stranded DNA genome of the Western Reserve (WR) strain of vaccinia encodes 229 genes (GeneBank accession n. NC_006998), whose transcription is regulated in a temporal manner (5,6). After entry into the cell, but before the release of viral DNA from the virion, early viral genes are transcribed by an encapsidated RNA transcription machinery (5). The resulting viral mRNA is released into the cytoplasm and translated by the host ribosomes. These early viral proteins suppress the hosts anti-viral response and allow the progression of vaccinia replication cycle, including the release of viral DNA from the virion (uncoating) (7,8). Once the vaccinia genome is released into the cytoplasm, viral DNA replication occurs in perinuclear structures known as viral factories (2). Once replication begins, transcription of post-replicative genes (intermediate and late genes) is initiated, leading to the production of proteins required for the assembly of new virions.

The cytosolic replication cycle, along with the presence of all essential proteins for DNA synthesis encoded within its genome, led to the belief that vaccinia DNA replication was independent of the host (9,10). However, more recent studies have shown that host proteins play a role in supporting viral replication (11-14). Moreover, several nuclear proteins involved in the maintenance of the host genome are recruited to cytosolic viral factories. Among these, topoisomerase 2α and β are the most abundant host proteins co-immunoprecipitating with nascent viral DNA (13). Unlike other viruses (15-18), vaccinia lacks a type II topoisomerase, suggesting that viral DNA replication relies on host topoisomerase 2.

Topoisomerase 2 (TOP2) plays a critical role in maintaining nuclear genome integrity by alleviating topological stress caused by the twisting and tangling of the DNA helix during DNA replication, transcription, and chromosome segregation (19). TOP2 resolves torsional strain by inducing transient double-strand breaks in the DNA and passing one strand through the break. This mechanism forms a temporary DNA-protein covalent complex, known as TOP2 cleavage complex, which is the target for several important anticancer drugs such as etoposide, doxorubicin, and mitoxantrone (20). Higher eukaryotes have two paralogs, TOP2α (TOP2A) and TOP2β (TOP2B) that are highly conserved in their core enzymatic domains, but diverge in their C-terminal regions (21,22). The divergent C-terminal domains (CTDs) are dispensable for their catalytic activity, but are extensively post translationally modified (https://www.phosphosite.org) and contribute to isoform-specific functions in vivo (22,23). TOP2A is highly expressed in proliferating cells where it plays a critical role in DNA replication and mitosis (24-26). In contrast, TOP2B is ubiquitously expressed throughout the cell cycle in both proliferating and differentiated cells where it is involved in transcriptional regulation and chromatin remodelling (27).

Previous studies have demonstrated that TOP2 poisons block vaccinia infection consistent with a possible role for the protein in vaccinia DNA replication (28,29). Interestingly, mutations in the vaccinia ligase gene (A50) were found to confer resistance to TOP2 poisons (28). This is because A50 recruits TOP2 to sites of viral DNA synthesis, thereby sensitizing the virus to TOP2-targeting drugs (30). Although the presence of TOP2A and TOP2B on cytosolic viral factories has been established (13,30), their specific roles during vaccinia replication and infection remain unexplored. We have now uncovered that TOP2 isoforms have non-redundant functions during vaccinia infection. TOP2A interacts with the viral DNA replication machinery to promote vaccinia DNA synthesis, while TOP2B contributes to antiviral RNA granule formation, supressing vaccinia replication.

## Methods

### Cell culture, infections, and drug treatments

HeLa and BS-C-1 cells were maintained in Dulbecco’s Modified Eagle Medium (DMEM) supplemented with 10% FBS, 100 U/ml penicillin, and 100 µg/ml streptomycin at 37 °C and 5% CO_2_. Vaccinia infections were performed as previously described (31) with wild type Western Reserve (WR) strain or the fluorescently tagged WR RFP-A3 strain (32) at an multiplicity of infection (MOI) =1, unless stated otherwise. ICRF-193 (I4659, Sigma, 7 μM) and Merbarone (M2070, Sigma, 80 μM) were used to inhibit the activity of TOP2. Viral protein synthesis and DNA replication were inhibited by cycloheximide (CHX, C7698, Sigma, 50 μM) and cytosine arabinoside (AraC, C1768, Sigma, 50 μM), respectively. Staurosporin (S4400, Sigma, 1 μM) was used to induce apoptosis. All drug treatments were performed throughout the time of infection.

### Generation of tagged constructs for expression during vaccinia infection

For transient expression during vaccinia infection, all open reading frames were placed under the control of the synthetic vaccinia early/late promoter (pE/L) (33). Full length TOP2A-GFP and TOP2B-GFP constructs were kindly provided by Dr. Christian Mielke, Heinrich Heine University, Duesseldorf (34). The original CMV promoter in these constructs was replaced by the pE/L promoter using Gibson Assembly (New England Biolabs), generating pEL-TOP2A-GFP and pEL-TOP2B-GFP. The pE/L promoter fragment used in the assembly was synthesized by Integrated DNA Technologies (IDT). To create pEL-TOP2A^ΔCTD^-GFP and pEL-TOP2B^ΔCTD^-GFP, deleted fragments in frame with GFP were synthesized by IDT and cloned into the PacI/ApaI or SnaBI/ApaI sites of pEL-TOP2A-GFP or pEL-TOP2B-GFP, respectively. TOP2A^CTD^ and TOP2B^CTD^ regions were amplified by PCR with primers containing MluI and ApaI restriction sites and subcloned into pEL-TOP2A-GFP and pEL-TOP2B-GFP, generating pEL-TOP2A^CTD^-GFP and pEL-TOP2B^CTD^-GFP. For the mutation of their nuclear localization signals (NLSs) (35) positively charged lysines (K) and arginines (R) were substituted with alanines (A). For TOP2A NLS, Ks and Rs within residues 1452–1469 and 1487–1494 were mutated (PAK*TK*NR*R*K*R*K*PS and TSK*K*SK*GE). Similarly, for TOP2B, residues within 1525–1545 and 1553–1569 were substituted (IPK*K*TTTPK*GK*GR*GAK*K*K*K*AS and PGR*K*TSK*TTSK*K*PK*K*TS). Mutated fragments were synthesized by IDT and cloned into the PacI/ApaI or Bsu36I/ApaI restriction sites of pEL-TOP2A^CTD^-GFP and pEL-TOP2B^CTD^-GFP using Gibson Assembly. The open reading frames of A20 and A50 were amplified from the vaccinia genome using primers containing EcoRI and NotI restriction sites and subcloned into pEL-GFP-N (36) and pEL-Flag-N, respectively.

### Transient transfections and knockdowns

The pE/L constructs (1.5 μg plasmid) were transfected into ∼10^6^ HeLa cells using 2 μl Lipofectamine 2000 (Thermofisher) at the time of infection. Protein expression was assessed 6-8 hours post infection (hpi). For knockdown experiments, HeLa cells were transfected with siRNA using Lipofectamine RNAi Max, according to the manufacturer’s protocol (Thermofisher). TOP2A and TOP2B knockdowns were achieved 72 h after transfection with 10 nM siRNAs (ON-TARGETplus SMARTpool, Dharmacon). For knockdown of viral proteins, 30 nM of siRNAs targeting D5 (5′-CGUAACACCUUGUGCAUUA[dT][dT],(8)), E3 (5′-AAGACUUAUGAUCCUCUCUCA [dT][dT]) or A50 (5′-GUGAAAGAGUACAAGUUCA[dT][dT],(8)) were transfected 24 h prior infection. Scrambled sequences were used as control at the same concentration as target oligos (AllStars Negative Control siRNA, Qiagen).

### Immunofluorescence and live imaging

Cells were plated on fibronectin-coated coverslips one day prior infection. At the indicated timepoints after infection, cells were fixed with 4% paraformaldehyde in PBS for 15 min, permeabilised with 0.1% Triton-X/PBS for 5 min and blocked in cytoskeletal buffer (1 mM MES, 15 mM NaCl, 0.5 mM EGTA, 0.5 mM MgCl_2_, and 0.5 mM glucose, pH 6.1) containing 2% (vol/vol) fetal calf serum and 1% (wt/vol) BSA for 30 min. Cells were stained with primary antibodies against TOP2A (Cell Signaling Technology, 12286, 1:250 dilution), TOP2B (Santa Cruz, sc-25330, 1:250), DDX1 (Proteintech, 11357-1-AP, 1:1,000 dilution), RTCB (Proteintech, 19809-1-AP, 1:50 dilution), G3BP1 (Proteintech, 66486-1-Ig, 1:1,000 dilution), FAM98A (Abcam, ab204083, 1:50 dilution), dsRNA (RNT-SCI-10010200, Scicons, 1:1,000 dilution), I3 (37) (1:1,000 dilution), followed by Alexa Fluor 488 or 568 conjugated secondary antibodies (Invitrogen; 1:500 in blocking buffer). Extracellular viruses were labelled with the 19C2 monoclonal against B5 (38) (1:1,000 dilution) followed by Alexa Fluor 647 anti-rat secondary antibody (Invitrogen; 1:500), before permeabilisation of the cells with detergent. Coverslips were stained with 4′,6-diamidino-2-phenylindole (DAPI) for 5 min, mounted on glass slides using Mowiol (Sigma), and imaged on a Zeiss Axioplan2 microscope equipped with a 63 x/1.4 NA Plan-Achromat objective and a Photometrics Cool Snap HQ cooled charge-coupled device camera. The microscope was controlled with MetaMorph 7.8.13.0 software. To achieve higher resolution, antiviral stress granules were imaged on an Olympus iX83 Microscope with Olympus 150x/1.45 NA X-Line Apochromatic Objective Lens, dual Photometrics BSI-Express sCMOS cameras, and CoolLED pE-300 Light Source (Visitech), which was controlled using Micro-Manager 2.0.0. For live imaging, cells were plated on fibronectin-coated iBidi chamber slides one day prior infection. Shortly before imaging, media was changed with Leibovitz’s L-15 Medium without phenol red (Thermofisher), supplemented with 10% FBS and the cell-permeant nuclear counterstain NucBlue™ Live ReadyProbes™ Reagent (Thermofisher). Cells were imaged on a Zeiss Axio Observer spinning-disk microscope equipped with a Plan Achromat 63×/1.46 NA oil lens, an Evolve 512 camera, and a Yokogawa CSUX spinning disk. The microscope was controlled by the SlideBook software (3i Intelligent Imaging Innovations). Quantification of B5 positive extracellular virus on the plasma membrane was performed using the analyse particle plugin in Fiji.

### Viral DNA and mRNA quantification

Viral DNA was quantified at 5 hpi, as this time point represents the peak of viral DNA synthesis (13). Total DNA was isolated from ∼10^6^ cells using DNeasy Blood and Tissue Kit, according to the manufacturer’s protocol (Qiagen), and quantified by spectrophotometry (Nanodrop, Thermofisher). Quantitative PCR was performed in triplicate on 384-well plates (Applied Biosystems) in 10 μl of total volume. For each reaction 20 ng of DNA template was amplified using PowerUp^™^ SYBR ^™^ Green Master Mix (Applied Biosystems) and 0.5 μM of primers specific to vaccinia genome. 18S rDNA was amplified as nuclear reference. Based on the temporal expression pattern of vaccinia genes (39), RNA levels of early genes were assessed at 2 hpi, while intermediate and late genes were evaluated at 5 hpi.Total RNA was extracted at these timepoints using RNeasy Mini Kit, according to the manufacturer’s protocol (Qiagen). One-step quantitative RT-PCR was performed on 50 ng RNA template using QuantiNova® SYBR® Green RT-PCR Kit (Qiagen), and 0.5 μM of primers specific to the indicated viral genes. GAPDH was used as housekeeping reference. The sequences of the primers used for quantitative PCRs are listed in Table S1. Changes in relative amount of viral DNA or mRNAs were calculated using the 2^-ΔΔCt^ method (40) and represented as fold changes relative to controls.

### Western Blotting

Cells were lysed in PBS supplemented with 1% SDS, protease/phosphatase inhibitor cocktail (Cell Signaling Technology), and Benzonase (Millipore). Proteins from the lysates were resolved on Bolt™ 4-12% Bis-Tris or NuPAGE™ 3-8% Tris-Acetate pre-cast gels (Thermofisher) and then transferred to nitrocellulose membranes. After blocking with 5% non-fat dry milk in PBS with 0.1% (v/v) Tween-20 (PBS-T) membranes were incubated overnight with primary antibodies against GFP (Clone 3E1, Crick Cell Services, 1:4,000 dilution), Flag (Sigma, F3165, 1:20,000 dilution), D5 ((41), 1:2,000 dilution), F12 ((42), 1:2,000 dilution), H5 ((43), 1:8,000 dilution), L4 ((44), 1:2,000 dilution), F13 ((45), 1:8,000 dilution), A27 ((46), 1:2,000 dilution), E9 ((47), 1:2,000 dilution), I3 ((30), 1:5,000 dilution), Vinculin (Sigma, V4505, 1:10,000 dilution), TOP2A (Invitrogen, MA5-12433, 1:2,000 dilution), TOP2B (Santa Cruz, sc-25330, 1:2,000 dilution), DDX1 (Proteintech, 11357-1-AP, 1:10,000 dilution), FAM98A (Abcam, ab204083, 1:2,000 dilution), RTCB (Proteintech, 19809-1-AP, 1:5,000 dilution), Actin B (Abcam, ab179467, 1:40,000 dilution), PARP (Cell Signaling Technology, #9542, 1:2,000 dilution). HRP conjugated secondary antibodies were obtained from Jackson ImmunoResearch and used at 1:10,000 in 5% milk in PBS-T. Immunoblots were developed using enhanced chemiluminescence (ECL) substrates (Thermofisher).

### Plaque assays and viral growth

Confluent monolayers of BS-C-1 cells were infected with vaccinia at a MOI of 0.1 in serum-free MEM. After one hour, the inoculum was replaced with a semi-solid overlay composed of a 1:1 mixture of MEM and 2% carboxymethyl cellulose, supplemented with the indicated concentrations of drugs or vehicle. At 72 hours post-infection, cells were fixed with 4% formaldehyde and stained for 30 minutes with crystal violet. Viral plaque-forming units per milliliter (PFU/ml) were quantified by counting the cleared plaques, while plaque size diameters was measured using the Fiji line tool. For quantification of viral growth, cells were infected in 6-well plates at an MOI of 1. At 8 hpi cells were scraped in the media and lysed by three freeze-and-thaw cycles. The amount of infectious virus produced was quantified by calculating the PFU/ml using plaque assays on confluent BS-C-1 cells.

### Co-Imminoprecipitation and Mass Spectrometry

HeLa cells were seeded in one or two 150 mm dishes per condition 24 hours before pull downs experiments to achieve approximately 80% confluency. Cells were simultaneously transfected with the pEL-GFP or Flag relevant constructs (25 μg DNA/ 150 mm dishes) and infected with vaccinia (MOI = 3). After 6 hours cells were collected in PBS and lysed in 10 mM Tris/HCl pH 7.5, 150 mM NaCl, 0.5 mM EDTA, 0.5 % Nonidet™ P40 Substitute, supplemented with Protease/Phosphatase Inhibitor Cocktail (Cell Signaling Technology). GFP was pulled down using GFP-Trap® Agarose as per manufacturer’s instructions (ChromoTek). Bead-bound protein samples were reduced, alkylated and digested with trypsin, essentially as previously described (48). The digests were subsequently analysed by LC-MS/MS on an Orbitrap Fusion Lumos, (ThermoScientific). All raw files were processed with MaxQuant 2.4.9.0 (49) using standard settings and searched against a UniProt Human Reviewed KB database. Statistical analysis was carried out using the Perseus module (v2.0.11) of MaxQuant. For Immunoblot analysis of co-immunoprecipitated proteins, beads were resuspended in 50 μl of Tris-Glycine SDS Sample Buffer (Thermofisher), boiled 10 minutes and loaded into gels for SDS-PAGE.

### dsRNA Slot Blot

Total RNA (4 μg) was diluted in RNAse free water to a final volume to 200 μl and transferred onto BrightStar™-Plus Positively Charged Nylon Membrane (Thermofisher) using a CSL Slot Blotting Unit (Cleaver Scientific LTD). RNA was UV crosslinked to the membrane with Stratagene stratalinker (1200 J/m2) and blocked with 5% non-fat dry milk in TBS with 0.1% (v/v) Tween-20 (TBS-T). A dsRNA antibody (RNT-SCI-10010200, Scicons, 1:2,000 dilution) was incubated for an hour in blocking buffer and detected with an HRP conjugated antibody as described in the Western blotting section. The dsRNA band intensity was quantified using Fiji.

### Statistical analysis and figure preparation

All data are presented as means ± S.E.M. For all experiments, means of at least three biological replicates were used to determine statistical significance by Dunnett’s multiple comparisons test. All data were analyzed using GraphPad Prism 10 and figures prepared using Illustrator 27.2.

## Results

### TOP2 cytosolic translocation and recruitment to viral factories are independent events

TOP2A and TOP2B are key regulators of genomic stability that are primarily localised in the nucleus of uninfected cells. Given this fact, we first investigated when and how they are recruited to cytosolic viral factories during vaccinia infection. TOP2s re-localisation occurred early as both proteins associated with cytoplasmic viral DNA by 2 hours post infection (Fig. 1A). At 7 hours post infection, TOP2A was located both, in the nucleus and at the viral factories, while TOP2B was completely recruited to viral factories (Fig. 1A). Given the rapid shutoff of host protein synthesis following vaccinia infection (50), we think it is more likely that the TOP2A recruited to viral factories originates from the nucleus, rather than from a newly synthesized cytoplasmic pool. TOP2A and TOP2B still translocated to the cytoplasm (diffuse cytosolic staining) when DNA replication, and therefore viral factory formation, was inhibited by AraC (Fig. 1B). This translocation was completely abrogated when viral protein synthesis was inhibited by Cycloheximide. Taken together, this suggests that the re-localisation of TOP2A and TOP2B to the cytosol requires early viral protein expression rather than the presence of viral DNA factories. To confirm this hypothesis, we inhibited viral uncoating, which prevents viral genome release, by knocking down the early viral protein D5 (Fig 1C) (8). In the absence of D5, TOP2A and TOP2B still translocated to the cytoplasm, confirming that their extranuclear localisation is triggered by early viral gene expression. Previous observations have demonstrated that the viral ligase A50, which is expressed early during infection, recruits TOP2 to viral factories (30). We therefore knocked down A50 to examine whether it is required for the cytoplasmic translocation of TOP2s. We found that, A50 was necessary to recruit TOP2A and TOP2B to viral factories but did not trigger their cytosolic translocation (Fig. 1E).

**Figure 1.**
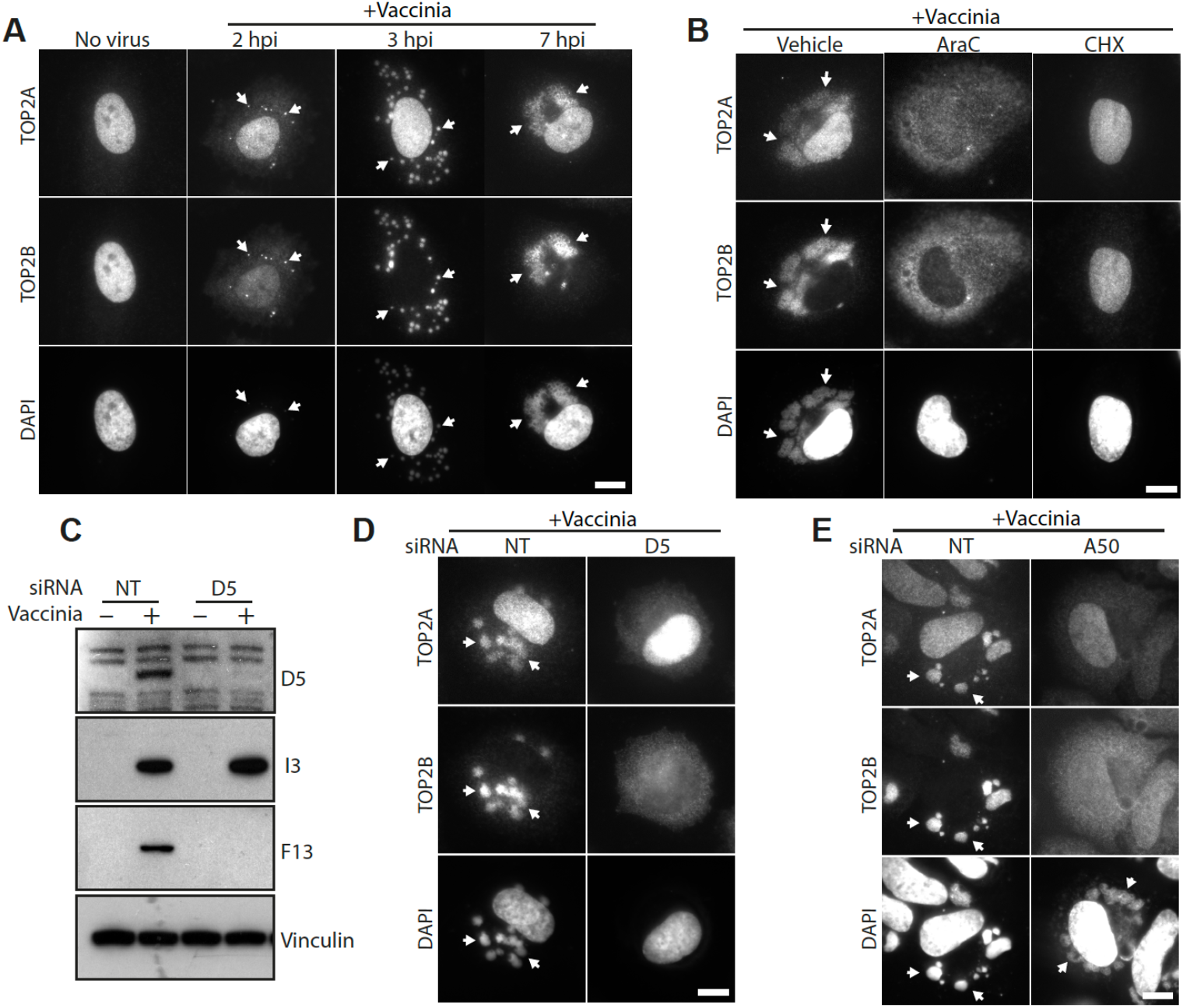
Cytosolic translocation of TOP2A and TOP2B during vaccinia infection. (**A**) Immunofluorescence analysis of HeLa cells infected with vaccinia virus for the indicated times demonstrates that endogenous TOP2A and TOP2B are recruited to cytosolic viral DNA early during infection (white arrows). (**B**) TOP2A and TOP2B translocation to the cytosol is dependent on early gene expression (Cycloheximide, CHX, treatment) but not viral DNA replication and factory formation (AraC treatment). (**C**) Immunoblot analysis reveals that knockdown of vaccinia uncoating factor D5 leads to loss of post-replicative (F13) but not early (I3) viral proteins. Vinculin represents the cell loading control. (**D**) Loss of D5 does not impair the cytosolic translocation of TOP2A and TOP2B, as both isoforms show diffuse cytosolic staining. (**E**) Knockdown of the viral ligase A50 prevents the recruitment of TOP2A and TOP2B to viral factories (white arrows) but does not affect their cytosolic translocation. Scale bars represent 10 µm.

### The C-terminal domains of TOP2 mediate the interaction with A50

To identify the regions of TOP2A and TOP2B responsible for their recruitment to viral DNA, we examined the localisation of a series of GFP tagged truncation mutants (Fig. 2A, Fig S1). Full-length TOP2A and TOP2B were recruited to both the nucleus and viral factories, whereas loss of their C-terminal domain (CTD) resulted in their cytosolic localization (Fig. 2B). This localisation is consistent with the fact that the CTDs contain the nuclear localization sequence (NLS) required for TOP2s nuclear import (35). Importantly, these C-terminally truncated proteins were not recruited to viral factories, suggesting that their CTDs mediate the interaction with A50. However, GFP-tagged CTDs did not associated with viral factories and were mostly nuclear. To abrogate the strong recruitment of the CTDs to the nucleus, we mutated one of the NLS sequences to weaken their nuclear targeting. NLS mutations impaired CTDs nuclear import, allowing recruitment of TOP2A and TOP2B CTDs from the cytoplasm to viral factories. GFP trap pulldowns on infected cells co-expressing GFP-tagged CTDs and Flag-tagged A50 demonstrated that the viral ligase associates with the CTDs of both isoforms, although it appears to have a greater preference for TOP2A (Fig. 2C). Together these results indicate that TOP2A and TOP2B are recruited to viral factories through an interaction between their CTDs and A50.

**Figure 2.**
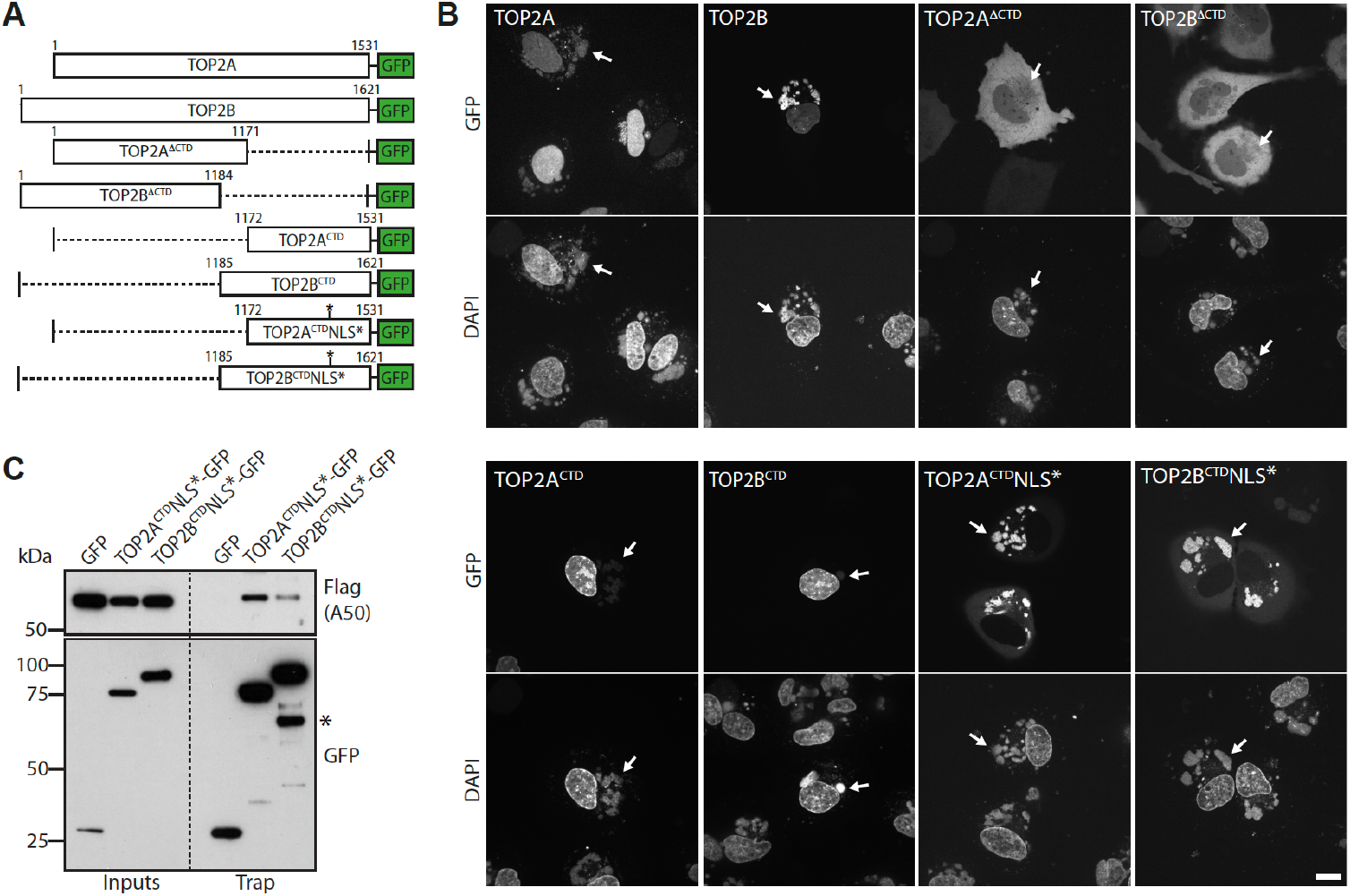
TOP2A and TOP2B interact with viral ligase A50 via their C-terminal domains. (**A**) Schematic representation of the GFP-tagged truncation mutants analysed in (B). (**B**) Confocal images showing the localisation of the indicated GFP-tagged TOP2A and TOP2B mutants in live infected HeLa cells at 6hpi. White arrows indicate cytosolic viral DNA. (**C**) Immunoblot analysis of GFP Trap pulldowns reveals that the TOP2A and TOP2B CTDs interact with Flag-tagged A50. * indicates a proteolytic product of TOP2B^CTD^NLS*-GFP. Scale bar represents 10 µm.

### Inhibition of TOP2 impairs vaccinia DNA replication and virion maturation

The recruitment of TOP2A and TOP2B to viral factories together with the observation that TOP2 poisons exhibit anti-poxviral activity (28,29), suggests a critical role for these enzymes in vaccinia DNA replication. To explore this hypothesis, we examined the impact of the TOP2 catalytic inhibitors merbarone and ICRF-193 on vaccinia infection. These inhibitors act at different stages of the TOP2 catalytic cycle: merbarone prevents the formation of the TOP2– DNA cleavage complex, while ICRF-193 stabilizes the noncovalent TOP2–DNA complex (20). Importantly, both inhibitors disrupt TOP2 function without trapping cleavage complexes, thereby causing significantly less DNA damage compared to TOP2 poisons. Both drugs inhibited plaque formation and reduced plaque size in a dose-dependent manner (Fig S2A, B and C). Viral DNA replication decreased by ∼90% in cells treated with either inhibitor compared to the DMSO control (Fig 3A). In line with this, the steady-state levels of post-replicative mRNAs and proteins were significantly reduced (Fig 3A, B). In contrast, the level of F12 (early viral protein) was unaffected even though the level of its mRNA was partially reduced by merbarone but not ICRF-193 (Fig 3A, B). Importantly, these effects were not due to cell death, as neither TOP2 inhibitor induced apoptosis during the experimental time course (8 hrs) (Fig. S2D). Consistent with the marked reduction in viral replication and post-replicative gene expression, virus assembly, as judged by cytoplasmic RFP-A3 positive virions and cell associated extracellular virions, was severely impaired in infected cells treated with the TOP2 inhibitors (Fig 3C, D).

**Figure 3.**
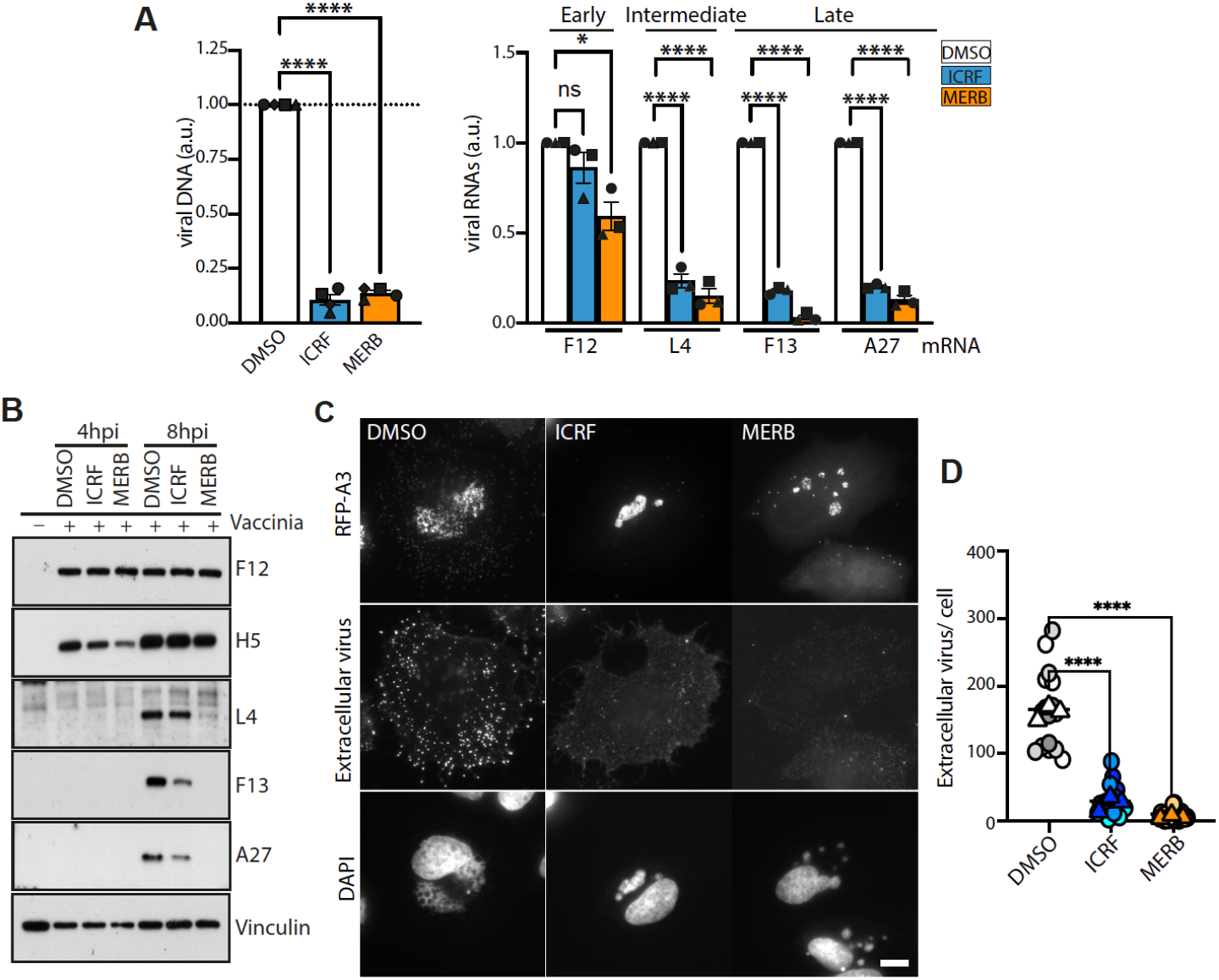
TOP2 inhibitors suppress vaccinia virus replication. (**A**) Treatment with the TOP2 inhibitors ICRF-193 and Merbarone significantly reduces viral DNA production (left) and steady-state levels of post-replicative intermediate (L4) and late (F13 and A27) viral transcripts (right). Data are presented as mean ± SEM from four and three biological replicates, respectively. Statistical significance was determined using one-way ANOVA followed by Dunnett’s post-hoc test for comparisons between treatment groups and the control. n.s. = not significant, *p ≤ 0.05, **p ≤ 0.01, ***p ≤ 0.001, ****p ≤ 0.0001. (**B**) Immunoblot analysis reveals that TOP2 inhibition reduces the expression of post-replicative intermediate (L4) and late (F13 and A27) but not early (F12) viral proteins. H5 is expressed both, early and late in infection. Vinculin represents the cell loading control. (**C**) Immunofluorescence analysis reveals that TOP2 inhibitors impair viral DNA factory formation and the assembly of new virions. RFP-A3 is a marker for virion cores (total virus). Extracellular virions associated with the plasma membrane during viral egress were detected using the B5 antibody. (**D**) Quantification of extracellular virions (B5 signal, as shown in panel C) at 8 hpi reveals a significant impairment in viral assembly and maturation in the presence of TOP2 inhibitors compared to the control. Drug treatments were maintained throughout the course of infection. Scale bar represents 10 µm.

### TOP2A and TOP2B have non-redundant functions during vaccinia replication

TOP2 inhibitors do not discriminate between the two isoforms as their catalytic domains have high degree of sequence homology. To determine whether TOP2A and TOP2B have distinct functions in vaccinia replication, we knocked down each isoform (Fig 4A). Interestingly, TOP2A knockdown led to a ∼50% reduction in viral DNA replication and post-replicative transcription, whereas TOP2B depletion enhanced these processes (Fig 4B, C). The steady-state levels of intermediate and late proteins also mirrored these trends (Fig 4D). Accordingly, virus assembly was impaired in the absence of TOP2A but not TOP2B (Fig 4E, F). Consistent with this observation, loss of TOP2A resulted in reduced virus production 8 hours post infection (Fig 4G). These results suggest that TOP2A and TOP2B have distinct roles in vaccinia replication.

**Figure 4.**
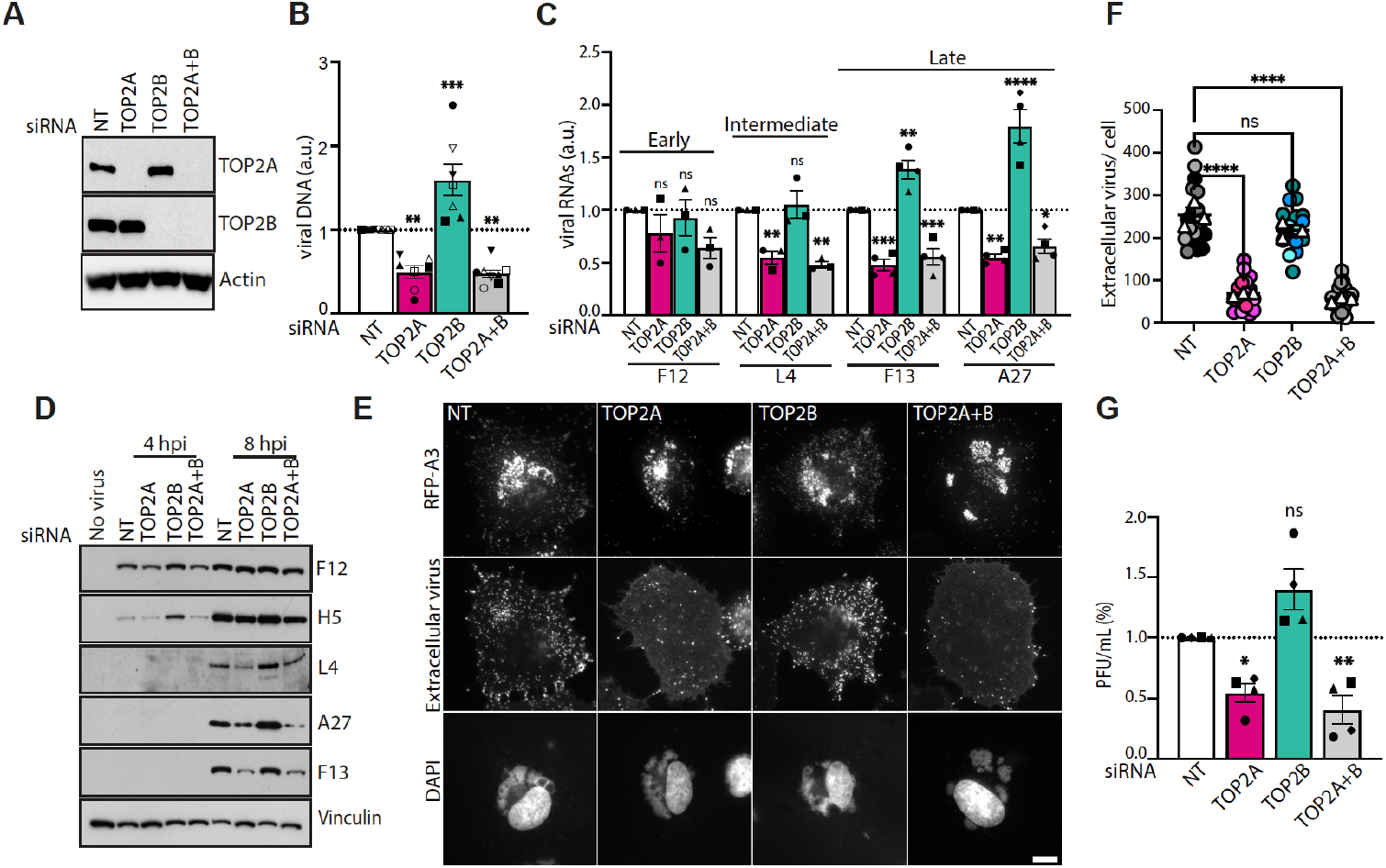
Opposing effects of TOP2A and TOP2B knockdowns on vaccinia replication. (**A**) Immunoblot analysis confirming siRNA-mediated knockdown of TOP2A and TOP2B. (**B**) Viral DNA production at 5 hours post-infection (hpi) is decreased by TOP2A knockdown and increased by TOP2B depletion. NT indicates non-targeting control siRNA. (**C**) The steady-state levels of intermediate (L4) and late (F13, and A27) viral transcripts are reduced and increased in TOP2A- and TOP2B depleted cells, respectively. (**D**) Immunoblot analysis reveals that loss of TOP2A and TOP2B results in reduced or increased expression of post-replicative viral proteins (L4, F13, and A27), respectively. (**E**) Immunofluorecence analysis reveals assembly of new virions at 8 hours post infection is severely impaired in TOP2A-depleted cells but unaffected by the loss of TOP2B. RFP-A3 is a marker for virion cores (total virus) and an antibody against B5 was used to detect extracellular virions associated with the plasma membrane during viral egress. Scale bar represents 10 µm. (**F**) Quantification of extracellular virions (B5 signal, as shown in panel E) reveals a significant impairment in viral assembly and maturation in the absence of TOP2A but not TOP2B. (**G**) Quantification of plaque-forming units (PFU) in cells depleted for TOP2A and/or TOP2B show a significantly impaired production of infectious virions in TOP2A-but not TOP2B depleted cells. All data in the bar chats are presented as mean ± SEM from at least three biological replicates. Statistical significance was determined using one-way ANOVA followed by Dunnett’s post-hoc test for comparisons between treatment groups and the NT control. n.s. = not significant, *p ≤ 0.05, **p ≤ 0.01, ***p ≤ 0.001, ****p ≤ 0.0001.

To uncover the mechanism behind this difference, we set out to identify isoform-specific interaction partners during vaccinia infection. TOP2 proteins are large and are only solubilised by relatively harsh cell lysis conditions (low levels of SDS) which are not conducive to co-immunoprecipitation of binding partners. We therefore chose to use the CTDs of TOP2A and TOP2B as baits because these domains mediate protein-protein interactions and confer isoform specificity in vivo (22,23). To identify infection-specific partners, we performed GFP pull downs on infected cells expressing GFP-tagged TOP2A or TOP2B CTDs with mutated NLS, as these proteins are recruited to viral factories, but not nuclear DNA (Fig. 2A, B). The volcano plot in Figure 5A shows the fold difference in the interaction partners of TOP2A and TOP2B CTDs during vaccinia infection determined by mass spectrometry. Most viral proteins interacted more strongly with TOP2A than TOP2B. Moreover, several of the TOP2A-enriched interacting proteins are involved in vaccinia DNA replication, including the viral DNA polymerase (E9), and Uracil-DNA glycosylase (D4). Only a few host proteins were found to interact preferentially with TOP2B during infection. Remarkably, four of these interactors (DDX1, FAM98A, RTCB, and C14ORF166) are part of the tRNA splicing ligase complex, which is involved in the formation and regulation of cytosolic RNA granules (51-55). Immunoblot analysis of GFP-trap pulldowns confirmed that TOP2A interacts with components of the viral DNA replication machinery E9, H5, and I3 (56-58) (Fig. 5B). These pulldowns also verified that TOP2B associates with DDX1, FAM98A, and RTCB, suggesting a potential role for this isoform in RNA metabolism during infection.

**Figure 5.**
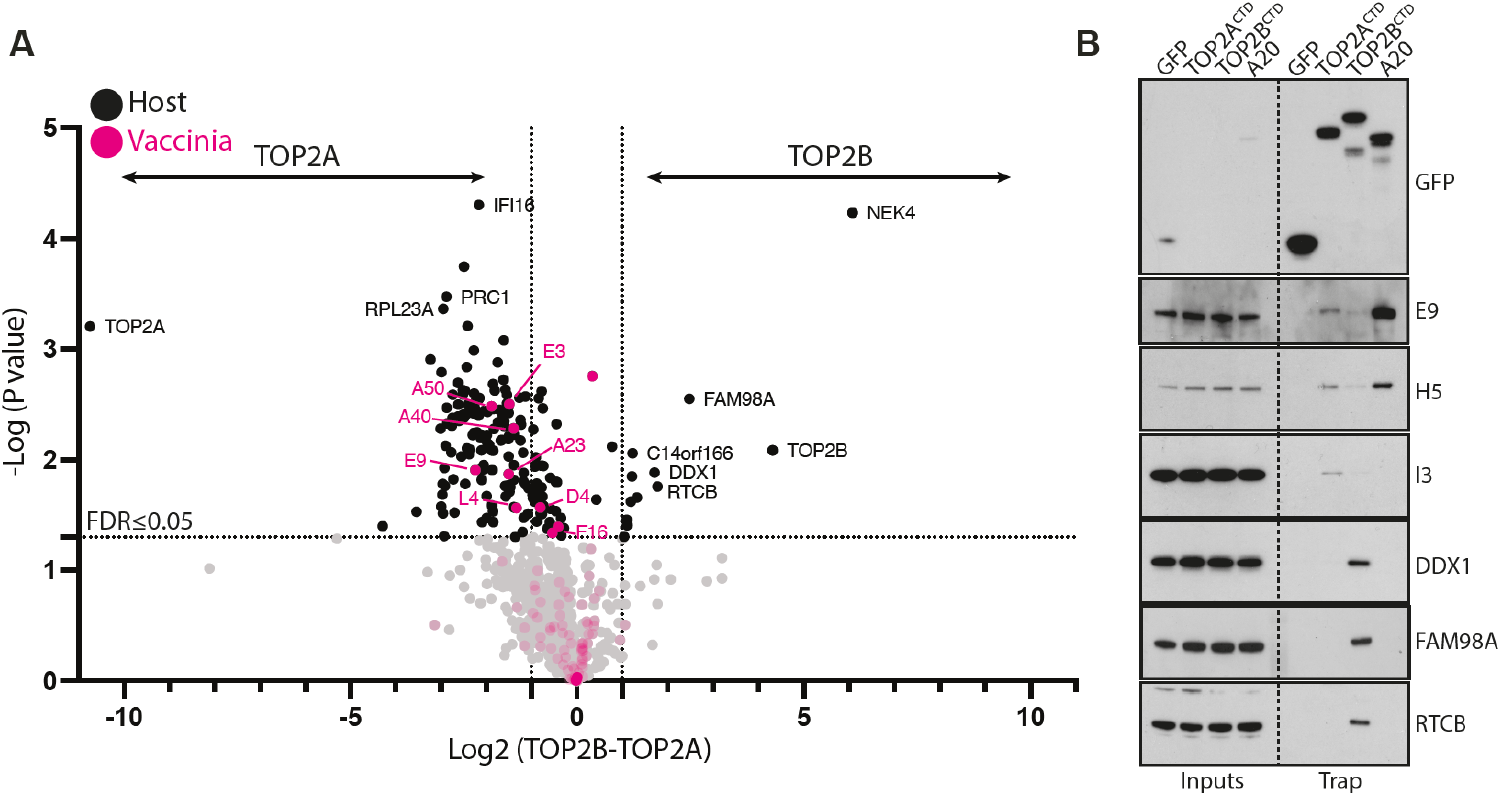
Identification of TOP2A and TOP2B interaction partners during infection. (**A**) The volcano plot shows proteins co-immunoprecipitating with TOP2A^CTD^NLS*-GFP or TOP2B^CTD^NLS*-GFP during infection identified by LC-MS in three technical replicates. Proteins significantly enriched (FDR≤0.05, at least two-fold) in TOP2A^CTD^ or TOP2B^CTD^ samples are shown in the upper left and upper right quadrants, respectively. Host and viral proteins are represented by black and magenta dots, respectively. (**B**) Immunoblot analysis of GFP-Trap pulldowns on infected cell lysates demonstrates that TOP2A^CTD^NLS*-GFP interacts with viral DNA replication proteins E9 (DNA polymerase), H5 (scaffold protein) and I3 (single-stranded DNA binding protein). In contrast, the TOP2B^CTD^NLS*-GFP interacts with components of the tRNA splicing ligase complex DDX1, RTCB, FAM98A, and C14orf166. A20, a component of the vaccinia DNA polymerase complex, served as a positive control and GFP alone represents a negative control. The GFP blot exposure was optimised for the Trap fractions. A longer exposure, optimised to show the input signals is shown in Fig.S3.

### TOP2B promotes dsRNA and antiviral granule formation

RNA granules form in the cytoplasm of eukaryotic cells in response to various stress conditions including viral infection (59). These stress granules consist of condensates of mRNA and RNA-binding proteins stalled in pre-initiation translation complexes. Stress granules-like structures also form during vaccinia infection as part of the host stress response (60-63). These specialized antiviral RNA granules (AVGs) form around replication factories when viral double-stranded RNA (dsRNA) is detected, inhibiting the production of viral proteins by blocking mRNA translation. However, vaccinia can evade this host defense, as the viral protein E3 binds and sequesters dsRNA, preventing AVG formation (64,65). Given that four of the identified TOP2B interaction partners play a role in RNA granule formation, we explored whether these proteins are components of AVGs during infection. To enhance the formation of AVGs, we silenced the expression of E3 and assessed the localization of DDX1, FAM98A, and RTCB at 6-8 hours post infection. We found that all three proteins co-localised with G3BP1, a well-established stress granule marker (66), in AVGs at the edge of viral factories (Fig. 6A). These AVGs also contained dsRNA as described previously (61) (Fig. 6B).

**Figure 6.**
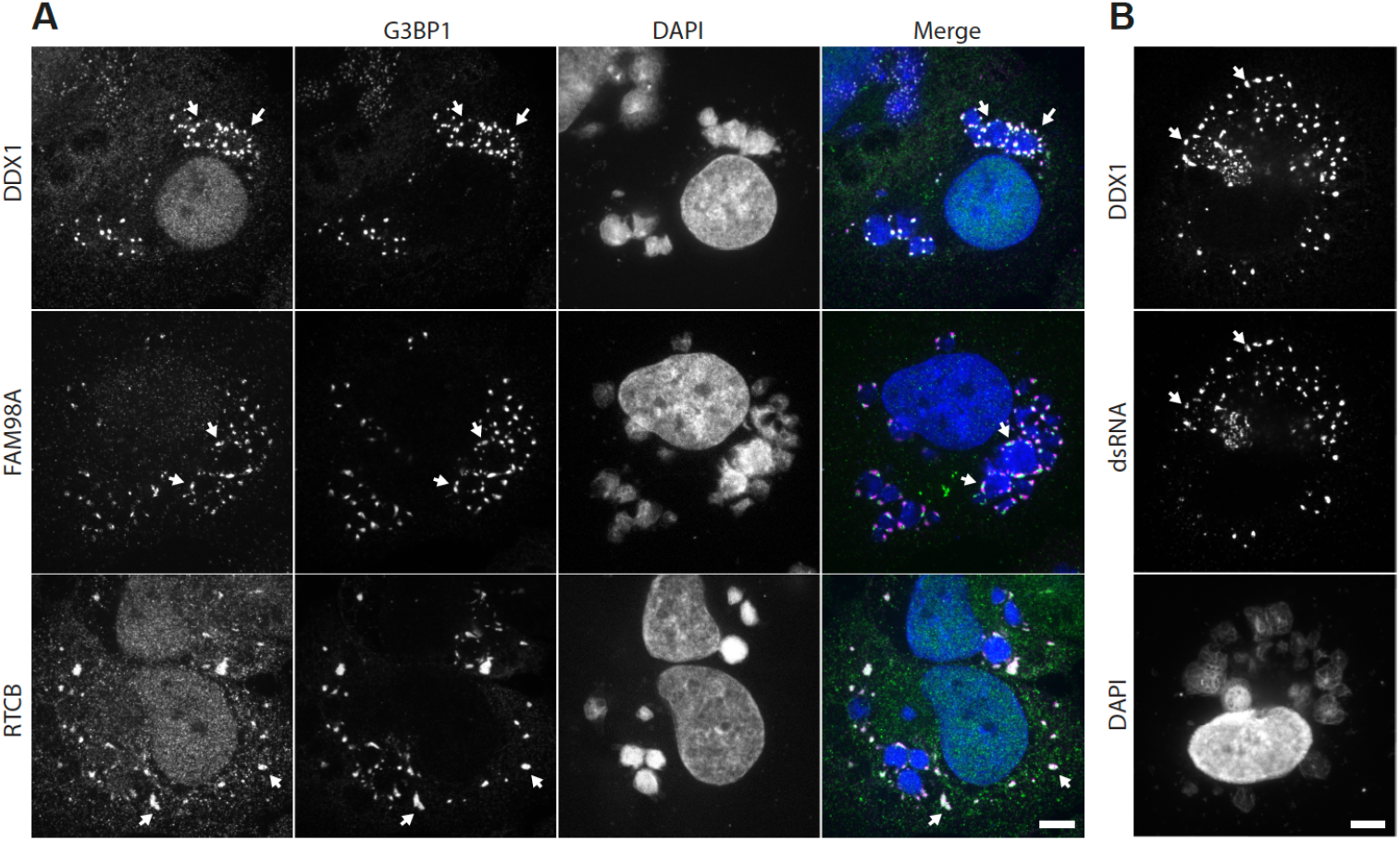
DDX1, FAM98A, and RTCB localize to antiviral stress granules. (**A**) Immunofluorescence images demonstrating that DDX1, FAM98A, and RTCB co-localize with G3BP1 in antiviral stress granules (AVGs – white arrows) around viral DNA factories 7 hours after infection in cells treated with E3 siRNA. (**B**) Immunofluorescence images show co-localization of double-stranded RNA (dsRNA) and DDX1 within AVGs. Scale bars represent 5 µm.

Having established that TOP2B interacts with components of AVGs, we investigated whether TOP2B contributes to their formation. To test this, we quantified the number of cells with AVGs during vaccinia infection in the presence or absence of TOP2B (Fig. 7A, B). We found that only ∼ 10% of infected cells had AVGs, which is consistent with previous reports (63). Only a small fraction of infected cells form AVGs because E3 binds to dsRNA suppressing their formation (64,67). Thus, E3 knockdown dramatically increased the percentage of cells with AVGs to ∼80%. We anticipated that any effect of TOP2B on AVG formation would be exacerbated in the absence of E3. Indeed, the percentage of cells with AVGs was significantly reduced in the absence of TOP2B (Fig. 7B). Depletion of TOP2B also strongly ameliorated the E3 knockdown phenotype, as there was a dramatic increase in expression of late viral proteins (Fig. 7C). This phenotype was specific for TOP2B, as TOP2A depletion had no effect on AVGs formation and did not increase late viral protein expression in the absence of E3 (Fig. S4). The loss of TOP2B but not TOP2A also resulted in a decrease in dsRNA formation during infection, which is consistent with a reduction in AVG formation (Fig. 7D). Taken together our observations indicate that TOP2B promotes AVGs formation, possibly suppressing viral replication, while TOP2A interacts with the viral replisome to enhance viral DNA replication.

**Figure 7.**
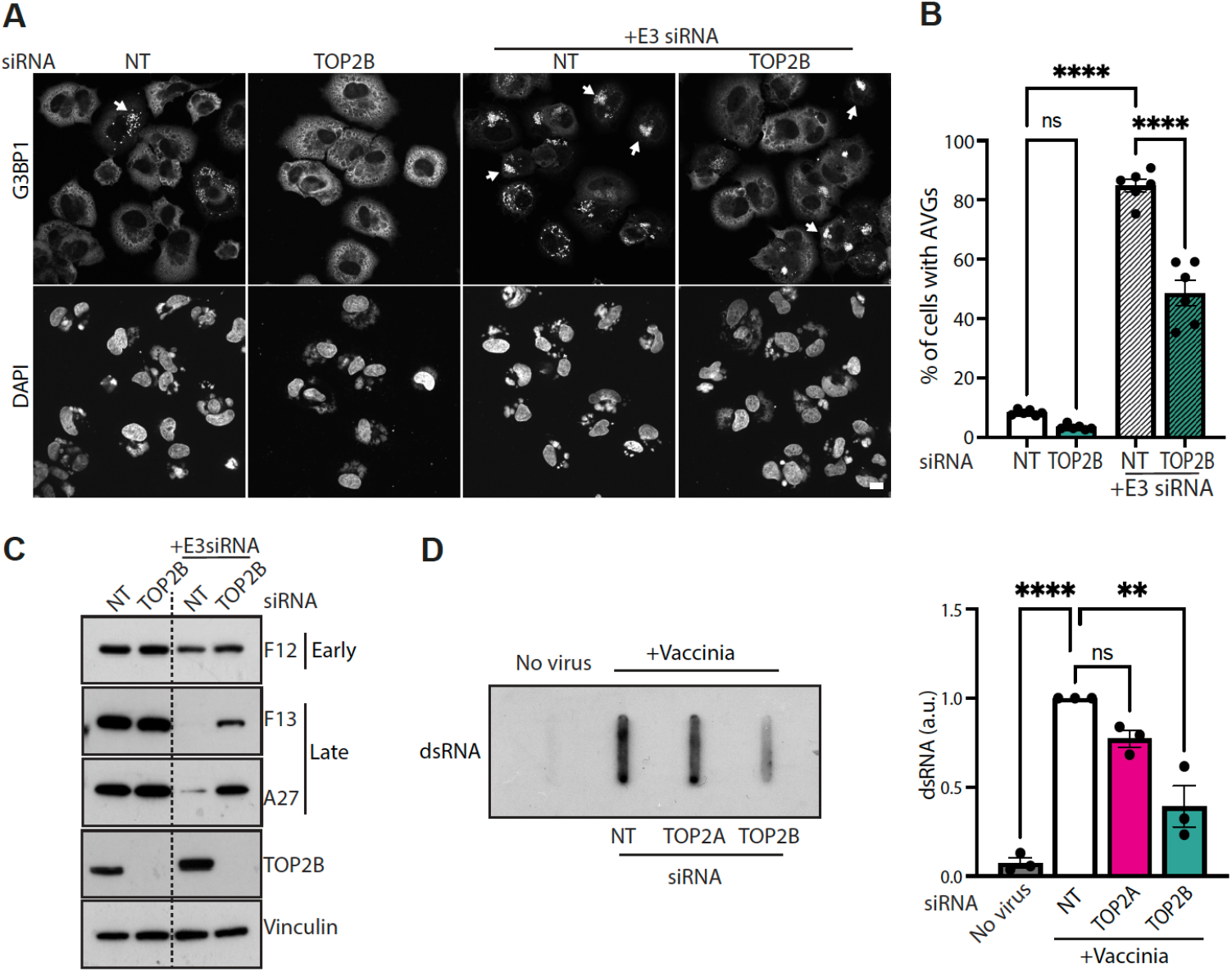
TOP2B promotes dsRNA production and antiviral stress granule formation. (**A**) Immunofluoresce images highlighting the presence of G3BP1 positive AVGs (white arrows) in vaccinia infected cells in the presence or absence of TOP2B and E3. Scale bar represents 10µm. (B) Quantification of the number of AVGs positive infected cells in the presence or absence of TOP2B and E3. Data represent mean ± SEM of counts from six biological replicates. Statistical significance was determined using one-way ANOVA followed by Dunnett’s post-hoc test for comparisons between treatment groups and the NT control; ****p ≤ 0.0001. (**C**) Immunoblot analysis reveals that loss of TOP2B and E3 results in increased expression of late viral proteins. Vinculin represents the cell loading control. (**D**) Slot blot analysis of dsRNA production in infected cells treated with TOP2B, TOP2A and non-targeting (NT) control siRNA (left) together with quantification of the levels of dsRNA (right). The data are from three biological replicates and error bars represent SEM. Statistical significance was determined using one-way ANOVA followed by Dunnett’s post-hoc test. n.s. = not significant, **p ≤ 0.01, ****p ≤ 0.0001.

## Discussion

Given previous observations showing TOP2A and TOP2B at sites of vaccinia DNA synthesis (13,30), we set out to better understand their role in viral replication. Our analysis reveals that vaccinia redirects TOP2A and TOP2B to cytosolic sites of viral replication early during infection in a two-step process. First TOP2s translocate from the nucleus to the cytosol and second, they are recruited to viral DNA through the interaction of their CTDs with A50, the viral ligase. The nuclear-to-cytosol translocation of TOP2s depends on early viral protein synthesis rather than being triggered by the presence of cytoplasmic viral DNA. This suggests that vaccinia has evolved mechanisms to actively recruit TOP2s from the nucleus to cytoplasmic viral factories to help facilitate DNA replication. The most obvious way this might be achieved is by modulating nuclear transport. Consistent with this notion, the nuclear pore complex has been shown to be necessary for viral morphogenesis (14). Further studies are required to identify the early viral protein(s) responsible for inducing TOP2 translocation and whether they also regulate the localisation of other nuclear proteins that have been identified at cytosolic sites of viral replication (11-13).

TOP2 poisons such as doxorubicin and mitoxantrone intercalate at the interface between DNA and the enzyme, impeding DNA religation and stabilizing the formation of TOP2–DNA cleavage complexes (20). This stabilization results in the accumulation of DNA double-strand breaks which activates DNA repair mechanisms (20,68). Given the extensive DNA damage and steric hindrance caused by TOP2–DNA cleavage complexes, it is not surprising that TOP2 poisons have a profound impact on vaccinia replication (28,29). In our study we evaluated the impact of ICRF-193 and merbarone on vaccinia replication as they inhibit the catalytic activity of TOP2 without inducing TOP2–DNA cleavage complexes (20). Both drugs also significantly affected vaccinia assembly by reducing viral DNA replication and post-replicative transcription by ∼90%. Isoform specific knockdowns allowed us to uncover that TOP2A and TOP2B have opposing roles during vaccinia replication. Depletion of TOP2A reduced vaccinia DNA replication and subsequent viral assembly by ∼50%, whereas TOP2B knockdown enhanced replication. Simultaneous knockdown of both isoforms mirrored the TOP2A phenotype, suggesting that TOP2B cannot functionally compensate for TOP2A. While TOP2A and TOP2B have nearly identical catalytic activities (69,70) and are potentially functionally redundant, they are known to perform distinct roles in nuclear genome maintenance (19,27), and this specificity is also evident in the context of vaccinia replication. Pharmacological inhibition of TOP2 had a significantly more pronounced effect on vaccinia replication compared to TOP2A knockdown. We believe this difference is explained by the fundamentally different approaches used to target the enzyme. In the case of TOP2A knockdown, we deplete the protein from the cell, whereas pharmacological inhibition blocks TOP2 activity by trapping both isoforms on the DNA. This trapping not only halts the enzymatic function of TOP2 but will also impede replication fork progression. Our data suggest that TOP2 inhibitors would have therapeutic potential against poxviral infections, with TOP2A specific inhibitors being particularly attractive candidates for antiviral treatment.

Our results demonstrate that TOP2A associates with components of the viral replisome to promote vaccinia DNA replication. Although the mechanism of vaccinia DNA replication still remains unclear, it is likely that topological stress accumulates as replication forks advance through constrained topological domains. The ability of TOP2A to alleviate such topological stress would undoubtedly improve replication efficiency. Furthermore, during vaccinia infection, DNA replication and post-replicative transcription occur simultaneously, potentially creating transcription-replication conflicts and helical tension that must be resolved by topoisomerases (71). Vaccinia encodes a type I topoisomerase that could partially address these issues, however, this enzyme primarily functions in early transcription and is not involved in DNA replication (72,73). The ability of TOP2A to promote replication suggests that vaccinia infection may proceed more efficiently in highly proliferative cells, where this isoform is highly expressed. It is possible that the elevated levels of TOP2A in cancer cells is one of the contributing factors in the preference of poxviruses for infecting tumours (74).

In contrast, TOP2B acts as a restriction factor during vaccinia infection, possibly by enhancing the antiviral response of the host. Our observation that TOP2B interacts with components regulating the formation of RNA granules provides insights into the antiviral properties of TOP2B (51-55). Stress granule-like antiviral granules (AVGs) are known to form at the periphery of viral factories during vaccinia infection (60-63). However, the TOP2B interactors we identified were not previously known to be associated with these structures. Outside of the context of vaccinia infection, DDX1, FAM98A, RTCB (also known as HSPC117), and C14ORF166 (also hCLE or RTRAF) are subunits of the pentameric human tRNA ligase complex, which mediates multiple RNA ligation reactions (55). This complex has been shown to play a role in promoting the replication cycle of human hepatitis delta virus, influenza and Sindbis virus (75-77). In contrast to its pro-replication role for these RNA viruses, our data suggest that the tRNA ligase complex restricts vaccinia replication. Understanding the precise role of this complex in AVG formation and vaccinia replication will provide new insights into host defence mechanisms during poxvirus infection.

The formation of AVGs is initiated by the detection of double-stranded RNA (dsRNA) by protein kinase R (PKR), which triggers the phosphorylation of the translation initiation factor eIF2α, leading to the arrest of viral protein synthesis (78). We suggest that TOP2B promotes AVG formation because it enhances dsRNA formation during transcription. In vaccinia infection, dsRNA is produced through the convergent transcription of genes on opposite DNA strands (5,79-81). Although vaccinia open reading frames typically do not overlap, post-replicative gene transcription lacks precise termination points, resulting in long run-on transcripts with heterogeneous 3’ ends (80-82). These transcripts can hybridize via complementary sequences in their 3’ ends to form dsRNA. TOP2B plays a critical role in nuclear transcription by regulating super helical tension generated by elongating RNA polymerase (38,83). By removing supercoils ahead of the transcription machinery, TOP2B facilitates transcription elongation, which is especially important for long and highly expressed genes (84,85). Based on this, we propose that TOP2B enhances the processivity of post-replicative vaccinia transcription, thereby generating longer run-on transcripts, resulting in more dsRNA. The increased production of dsRNA associated with TOP2B triggers antiviral defences, suggesting that its recruitment may be an unintended consequence of a strategy to recruit TOP2A to viral factories. It is also interesting that the viral ligase A50 has a stronger preference for TOP2A compared to TOP2B. The recruitment of TOP2B, while deleterious to viral replication, is thus tolerated as a trade-off that is less detrimental than not recruiting TOP2A.

In summary, we have demonstrated that while the mechanism underlying TOP2A and TOP2B recruitment to sites of viral replication is shared, the two isoforms perform distinct functions on vaccinia DNA that lead to opposite outcomes. Moreover, vaccinia infection now provides a unique model for studying mechanisms regulating the cellular localisation and differential functions of topoisomerase 2 α and β.

## Supporting information

Combined supplemental figures and legends

## Data Availability

The data underlying this article will be shared on request to the corresponding author.

## Supplementary Data Statement

Supplementary Data are available at NAR Online

## Acknowledgments

We thank Mark Skehel in the Crick proteomics science technology platform for performing mass spec analysis, Christian Mielke, Heinrich Heine University, Duesseldorf for providing GFP-tagged TOP2A and TOP2B constructs as well as Gabrielle Larocque and Antonio Postigo for generating TOP2A^ΔCTD^-GFP and GFP-A20 clones respectively. We also thank Angika Basant (King’s College London) as well as Simon Boulton, Snezhka Oliferenko, Frank Uhlmann and Miguel Hernandez Gonzalez at the Francis Crick institute for feedback on the manuscript. For the purpose of open access, the authors have applied a CC BY public copyright licence to any author accepted manuscript version arising from this submission.

## Author Contributions

**IDR:** Conceptualization, formal analysis, investigation, writing - original draft, writing – review & editing. **LK:** formal analysis, investigation. **MW:** Conceptualization, funding acquisition, supervision, writing – review & editing.

## Funding

This work was supported by the Francis Crick Institute, which receives its core funding from Cancer Research UK [CC2096], the UK Medical Research Council [CC2096], and the Wellcome Trust [CC2096].

## Conflict of interest disclosure

The authors declare there are no conflicts of interest.

## Notes

### Competing Interest Statement

The authors have declared no competing interest.

### Summary of Updates

One additional experiment has been performed and the text has been substantially revised to address the 4 reviewers comments.

