## Supplementary material for "Non-Redundant Roles of Topoisomerase 2α and 2β in the Cytosolic Replication of Vaccinia Virus": Combined supplemental figures and legends

**Supplemental figures - Dalla Rosa et al.**

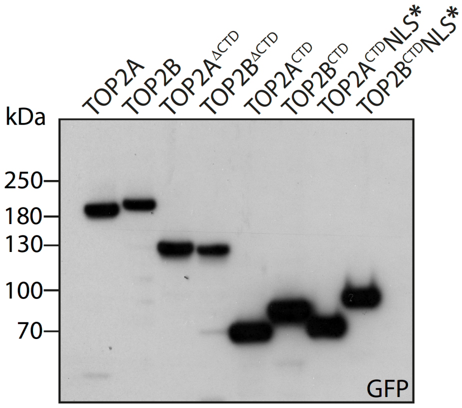


**Figure S1. Expression of GFP-tagged TOP2A and TOP2B mutants.** Immunoblot analysis with anti-GFP antibody confirms all GFP-tagged TOP2A and TOP2B mutants are expressed at their expected molecular weights.

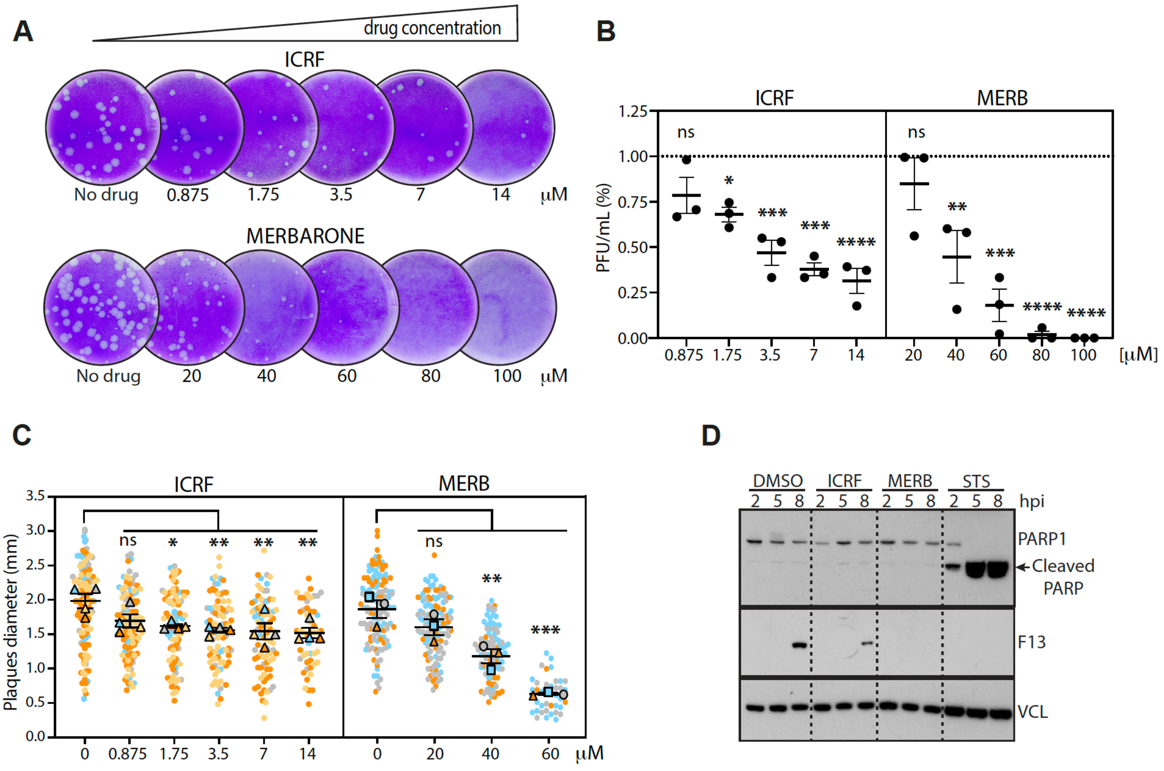


**Figure S2. ICRF-193 and Merbarone inhibit vaccinia plaque formation.**
(**A**) Representative images of plaque assays performed with increasing concentrations of ICRF-193 or Merbarone. Quantification of plaque numbers reveals a dose-dependent reduction in plaque formation (**B**) and plaque size (**C**) in the presence of TOP2s inhibitors. Data are presented as mean ± SEM from three biological replicates. Statistical significance was determined using one-way ANOVA followed by Dunnett's post-hoc test for comparisons between treatment groups and the control. n.s. = no significance, $*p \leq 0.05$, $**p \leq 0.01$, $***p \leq 0.001$, $****p \leq 0.0001$. (**D**) Western blot analysis of PARP cleavage shows that ICRF-193 or Merbarone treatments do not cause apoptosis of infected cells within 8 hours treatment. Staurosporin is used as positive control.

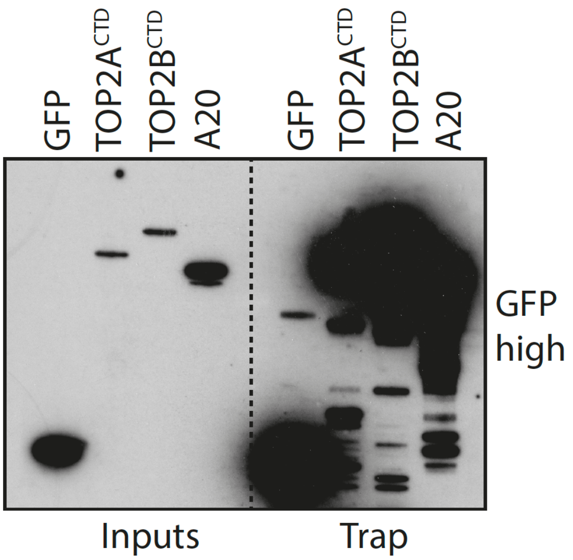


**Figure S3. Immunoblot analysis of GFP-Trap pulldowns.** A longer exposure of the GFP blot in Fig. 5B shows the expression levels of GFP-tagged constructs in the input samples.

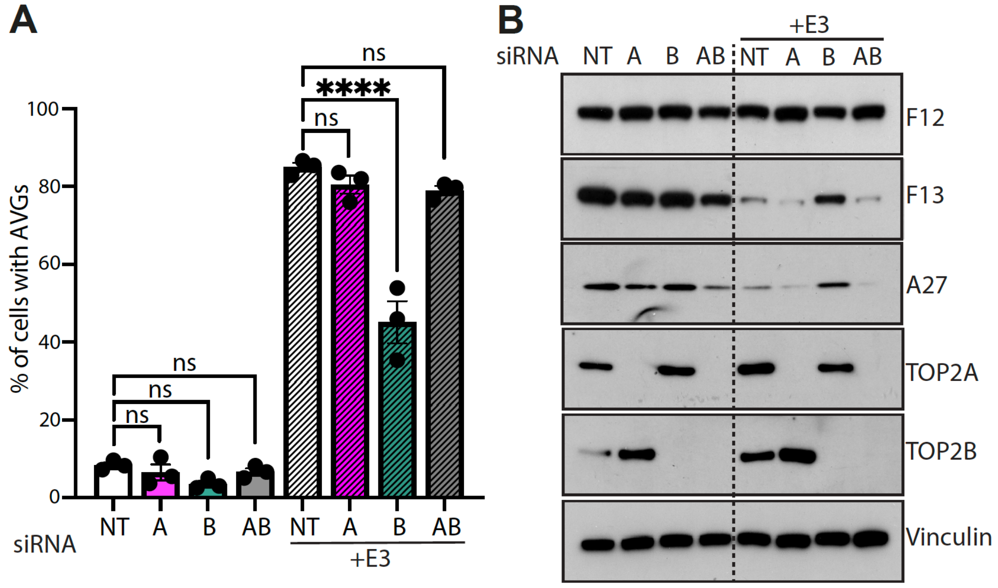


**Figure S4. TOP2A knockdown does not affect formation of antiviral stress granules.** (A) Quantification of AVG formation in the presence or absence of TOP2A or TOP2B. Data are presented as mean ± SEM from three biological replicates. Statistical significance was assessed using one-way ANOVA followed by Dunnett's post-hoc test for comparisons between treatment groups and non-targeting (NT) controls. n.s. = not significant, ****$p \leq 0.0001$. (B) immunoblot analysis of the indicated proteins reveals knockdown of TOP2B but not TOP2A restores the expression of late viral proteins in infected cells in the absence of E3. NT represents non-targeting siRNA control and vinculin is the cell loading control.

**Table S1: List of primers used for qPCR**

| | Sequence (5'-3') |
|---|---|
| 18S DNA Fw | GTCCAAAGCAGGCCCGAG |
| 18S DNA Rev | CCCTCTTAATCATGGCCTC |
| VACV DNA Fw | CCACGTTTGTTCATATACTAC |
| VACV DNA Rev | GTAAGAAGTAAATGCGTGC |
| GAPDH mRNA Fw | ACAGTTGCCATGTAGACC |
| GAPDH mRNA Rev | TTTTTGGTTGAGCACAGG |
| F12 mRNA Fw | GTTTGATCTAACGAGGAAG |
| F12 mRNA Rev | GGGCATATAGGTCATCATT |
| L4 mRNA Fw | CACAGGAGCAATGTTTACC |
| L4 mRNA Rev | GACGGCAGTCATATCCGTA |
| F13 mRNA Fw | GAACATATTCGTCGTCGAC |
| F13 mRNA Rev | CTAGTTGGTAATTGGGATAAG |
| A27 mRNA Fw | CTCTTAGAGTTTCAGCGTG |
| A27 mRNA Rev | GACGACAATGAGGAAACTC |
